# Mechanosensitive channel-based optical membrane tension biosensor

**DOI:** 10.1101/2022.10.04.510821

**Authors:** Yen-Yu Hsu, Agnes M. Resto Irizarry, Jianping Fu, Allen P. Liu

## Abstract

Plasma membrane tension functions as a global physical organizer of cellular activities. Technical limitations of current membrane tension measurement techniques have hampered in-depth investigation of cellular membrane biophysics and the role of plasma membrane tension in regulating cellular processes. Here, we develop an optical membrane tension biosensor by repurposing an *E. coli* mechanosensitive channel via insertion of circularly permuted GFP (cpGFP), which undergoes a large conformational rearrangement associated with channel activation and thus fluorescence intensity changes under increased membrane tension.

---

The plasma membrane is the initial medium through which a cell detects and reacts to external physical stimuli. As a global mechanical regulator, plasma membrane tension coordinates a number of cellular behaviors^1–3^. For instance, membrane tension governs the homeostasis between exocytosis and endocytosis, where high membrane tension promotes exocytosis while low membrane tension supports endocytosis by enhancing vesicle trafficking^4,5^. In addition, actin-based protrusions have also been suggested to be regulated by plasma membrane tension wherein low membrane tension facilitates polymerization of actin filaments, which generates lamellipodium-like protrusions at the leading edge during cell spreading^6^. Moreover, plasma membrane coordinates cytoskeletal tension for force transmission to control cell polarity and movement^7^.

Although membrane tension has been shown to play critical roles in regulating cell behaviors, the molecular mechanisms by which cells actively regulate membrane tension remain largely unknown. Studies have shown that certain membrane proteins can sense extracellular mechanical forces and convert them into intracellular biochemical signals dubbed as mechanotransduction to modulate cell activities. For example, changes in membrane tension can be recognized and be affected by the activity of membrane-to-cortex proteins, which act as a linker between cellular membranes and actin filaments^8^. Mechanosensitive (MS) channels are another membrane tension sensor with their channel pore opening controlled by changes in plasma membrane tension^9^. Found in all bacteria, mechanosensitive channel of large conductance (MscL) is one of the most studied non-selective MS channels for transporting molecules and ions across the membrane in response to physical stimuli^10^. It is also believed to function as a pressure regulator to prevent lysis of bacterial membrane resulting from the sudden increase of membrane tension during osmotic down-shock^11^. Other eukaryotic MS proteins such as transient receptor potential (TRP) channels^12^ and Piezo1^13^ are found to mediate a variety of sensations including hearing, touch, and thermo- and osmo-sensation.

Despite the above-mentioned proteins identified as natural membrane tension sensors, a lack of suitable membrane tension measurement tools has hindered detailed studies of the effect of membrane tension on cellular processes. Therefore, development of molecular sensors that report membrane tension of live cells is of significant interest. One of the most widely used technologies for membrane tension quantification is micropipette aspiration^14^, in which surface tension is deduced by measured radius of pipette and cells subjected to suction pressure applied by the micropipette using the Laplace law. The use of optical tweezers^15^ or atomic force microscopy^16^ is another option for indirect membrane tension measurement. However, these methods require special technical expertise and instrumentation. In recent years, several new methods have been developed to measure molecular tension. The development of fluorescence resonance energy transfer (FRET)-based biosensors has revolutionized the imaging of molecular signals with high spatiotemporal resolution^17,18^. However, the limitations in sensitivity and specificity have hindered their broader applications. In addition, current FRET-based tension biosensors focus on the tension change in the actin cytoskeleton or focal adhesion proteins rather than plasma membrane tension^19^. Although fluorescent membrane tension probes using small molecules such as Laurdan and Flipper-TR have been used in the past few years^20,21^, their functions and sensitivities are highly dependent on lipid and chemical environment. To overcome above-mentioned technical barriers, herein we report the development of an optical membrane tension biosensor using genetically modified MscL.

Specifically, our approach involves the development of a membrane tension OFF senor, by inserting circularly permuted green fluorescence protein (cpGFP), whose optical property depends on its structure, into the extracellular domain of MscL G22S. As a gain-of-function mutant of MscL, MscL G22S has a ~50% lower tension activation threshold (5-6 mN/m) compared to wild type (WT) MscL^22^. MscL G22S is used instead of WT MscL given the increased sensor sensitivity to mechanical stimuli. An increase in membrane tension causes the tilting of MscL G22S’s transmembrane helices and triggers channel opening. We hypothesize that structural rearrangement of MscL G22S caused by membrane tension would be transferred to the connected cpGFP, leading to a conformational change that gives rise to a change in its fluorescence (Figure 1A). A similar concept has been recently utilized for the successful development of a fluorescent voltage sensor^23^.

**Figure 1.**
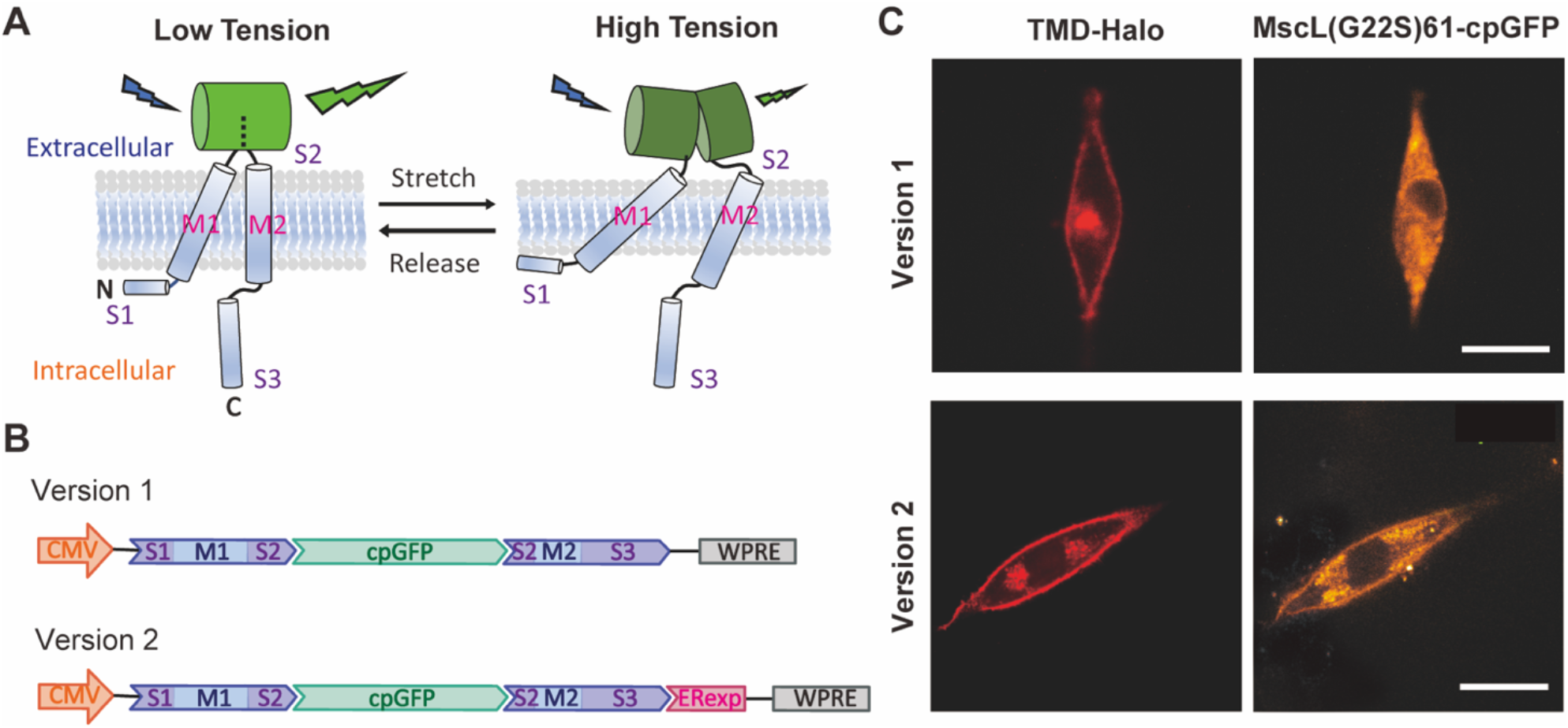
Schematics of the structure of the MscL membrane tension biosensor and its expression in living cells. (A) The schematic of engineering MscL as a membrane tension sensor using cpGFP. The substantial conformational changes of the extracellular domain S2 caused by the tilting of the transmembrane domain M1 and M2 under high tension leads to a decrease in fluorescence while cpGFP stays fluorescent under low tension. For simplicity, only a monomer is illustrated with S1, M1, S2, M2, and S3 domains shown. (B) The DNA constructs of the version 1 (top) and version 2 (bottom) MscL membrane tension biosensors. For both constructs, cpGFP was inserted to the sensory domain of MscL G22S after amino acid 61 as an OFF sensor. Version 2 is optimized with the addition of Kir2.1 ER export signal (ERexp) for enhancement of membrane localization. (C) Confocal fluorescence images showing the expression of version 1 (top) and version 2 (bottom) constructs in NIH3T3 cells. Scale bar: 5 μm.

To generate the tension biosensor DNA construct, we first encoded MscL G22S with cpGFP fused to its extracellular loop domain after amino acid 61 into a lentivirus vector, pLVX-Puro vector, under the control of CMV promoter as version 1 (Figure 1B). However, transfection of NIH3T3 cells leads to expression of the sensor version 1 and their notable intracellular accumulation but poor membrane localization (Figure 1C). We speculate a lack of a mammalian-specific export signal, which inhibits heterologous protein retention in the endoplasmic reticulum (ER), is responsible for MscL cytoplasmic accumulation. To overcome this, we next appended Kir2.1 ER export signal (FCYENEV), which is known to mediate protein export from ER to Golgi complex to plasma membrane^24^, to the C terminus of MscL G22S as version 2 (Figure 1B). This strategy has previously been shown to promote MscL membrane expression in neurons^25^. To examine localization of the MscL tension biosensors we co-transfected cells with either biosensor design along with a membrane-targeted HaloTag construct (TMD-Halo), where HaloTag was linked to the transmembrane domain of transferrin receptor (Tf R) for membrane localization.

As shown in Figure 1C, reduction of cytoplasmic aggregation and enhancement of membrane expression of version 2 biosensor was confirmed by stronger cpGFP signal observed in plasma membrane targeted with TMD-Halo as compared to that of version 1 biosensor. Version 2 biosensor (MscL(G22S)61-cpGFP) is referred to as MscL tension biosensor and used for the remainder experiments.

To test MscL tension biosensor sensitivity, we applied an osmotic pressure test, with increasing osmotic shock to cells transfected with the MscL tension biosensor. Previously, we have shown influx of small molecules into mammalian cells expressing MscL in an osmotic pressure dependent manner^26,27^. In the osmotic pressure test, the osmolarity of culture medium was gradually decreased from 330 to 108 mOsm. As shown in Figure 2Ai, MscL tension biosensors were found on the membrane as indicated by TMD-Halo in NIH3T3 cells over the range of hypo-osmotic conditions. MscL(G22S)61-cpGFP fluorescence decreased as the osmolarity of culture medium decreased, supporting that the MscL tension sensors function properly in living cells.

**Figure 2.**
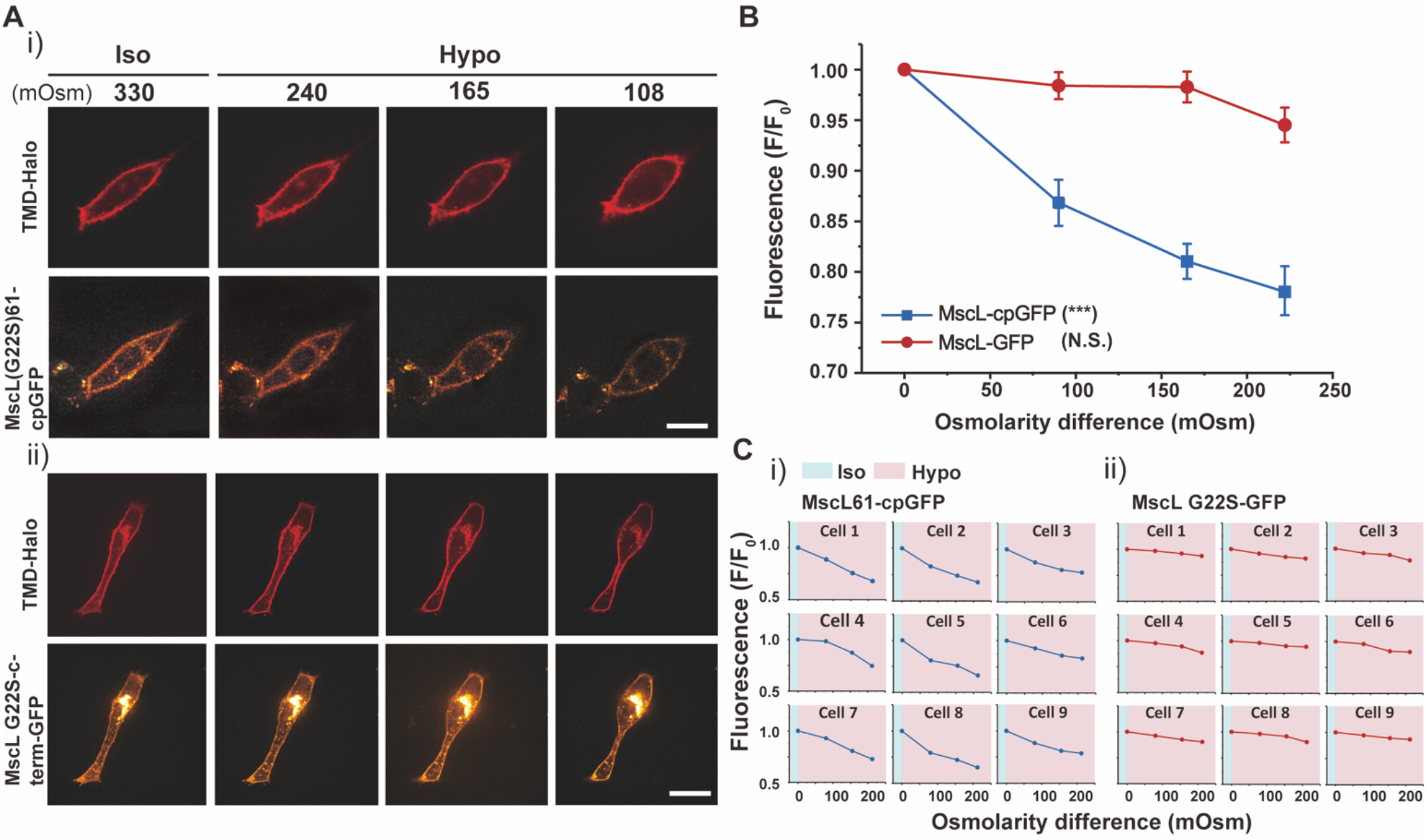
NIH3T3 cells transfected with the MscL tension biosensor in response to osmotic pressure. (A) (i) Confocal images showing the membrane localization of the MscL(G22S)61-cpGFP in NIH3T3 cells in response to different osmotic shocks. Transfected cells were cultured in iso-osmotic condition for 2 days and then DI water was sequentially added to the cell culture media to create increasing hypo-osmotic environments. Each image was taken four minutes after the addition of DI water. (ii) Confocal images showing the NIH3T3 cells transfected with the MscL(G22S)-c-term-GFP construct as a control in response to different osmotic shocks. The experiment followed the same method as mentioned in (i). (B) Normalized fluorescence intensities of the cell membranes of NIH3T3 cells transfected with the MscL tension biosensor (MscL(G22S)61-cpGFP) or MscL(G22S)-c-term-GFP under different osmotic conditions corresponding to the experiment mentioned in (A). (C) Normalized fluorescence intensities of the MscL tension biosensor (i) and MscL(G22S)-c-term-GFP control (ii) localizing on cell membranes of nine different NIH3T3 cells, from three separate experiments, at increasing osmotic pressures. Scale bars: 10 μm. The error bars denote standard error. ***: *p* < 0.001.

To verify that the declining fluorescence intensity in Figure 2Ai was not caused mainly by photobleaching, we fused GFP to the C-terminus of MscL G22S as a control (MscL(G22S)-c-term-GFP) and carried out the same osmotic pressure test. While the membrane localization of the control sensor was similar to that of the MscL tension biosensor, no significant change of the green fluorescence intensity at the plasma membrane was detected as the osmolarity of culture medium decreased (Figure 2Aii). We analyzed the fluorescence intensity by normalizing the fluorescence with the initial fluorescence intensity localized to the plasma membranes (using a mask generated from the TMD-Halo image) of cells under iso-osmotic condition (Figure S1). Cells with MscL tension biosensors displayed a substantial decrease in fluorescence in response to increasing osmotic pressure. For example, osmotic pressure with an osmolarity difference of 220 mOsm reduced fluorescence intensity by about 23%, whereas cells expressing the control sensors manifested only 5% fluorescence reductions under the same osmolarity difference (Figure 2B). This is also evident from normalized fluorescence of MscL tension biosensors in individual cells for both conditions (Figure 2C). Besides MscL(G22S)-c-term-GFP, we also generated three other controls. The first construct is GFP-P2A-MscL(G22S), in which we used a bicistronic construct with self-cleaving peptide P2A to express MscL G22S and GFP. To rule out the influence of different environmental sensitivity and photostability between GFP and cpGFP on fluorescence change in response to osmolarity, we created another control MscL G22S-c-term-cpGFP, where cpGFP is fused to the C-terminus of MscL G22S. In addition, we also inserted cpGFP into the extracellular domain after amino acid 70 of a glutamine transporter GluR0, an ion channel which is not mechanosensitive, as Glu70-cpGFP. With all three control constructs, we found that GFP/cpGFP fluorescence at the membrane (GFP-P2A-MscL(G22S), MscL(G22S)-c-term-cpGFP and Glu70-cpGFP) did not reduce with increasing osmolarity difference (Figure S2-S4), suggesting that volume changes of cells due to osmotic downshock did not appreciably change fluorescence of these control constructs at the membrane.

Besides testing hypotonic conditions, we also applied increasing hyper-osmotic shocks to cells by adding different amount of 10X PBS to the cell culture medium and found that no appreciable cpGFP membrane fluorescence changes were observed during the test (Figure S5), consistent with the fact that hypertonic state does not change the protein structure of MscL, and MscL remains closed under hyper-osmotic conditions as in iso-osmotic condition.

To further verify the sensitivity of fluorescence shift of the MscL tension biosensor in response to changes in membrane tension, we performed cyclic osmotic test on NIH3T3 cells co-transfected with MscL tension biosensor and TMD-Halo. The osmolarity was repeatedly switched between iso-osmotic (~330 mOsm) and hypo-osmotic conditions (~165 mOsm). As shown in Figure 3A, MscL tension biosensor was located on plasma membrane as expected. MscL(G22S)61-cpGFP fluorescence intensity varied depending on membrane tensions resulting from the cyclic iso-osmotic and hypo-osmotic conditions. We observed an alternating pattern of high and low fluorescence in iso-osmotic and hypo-osmotic conditions, respectively. When quantified as before, fluorescence intensities of the MscL tension biosensor displayed a ‘W’ shape trend (Figure 3B, Ci). A similar cyclical test was also conducted on NIH 3T3 cells transfected with MscL-(G22S)-c-term-GFP as a control, in which no obvious trend of MscL fluorescence could be correlated to the osmolarity changes (Figure 3Aii, B, Cii). Our other controls GFP-P2A-MscL(G22S), MscL(G22S)-c-term-cpGFP and Glu70-cpGFP also lacked such a trend (Figure S6-S8). Together, these results demonstrate the reversibility of the MscL tension biosensor.

**Figure 3.**
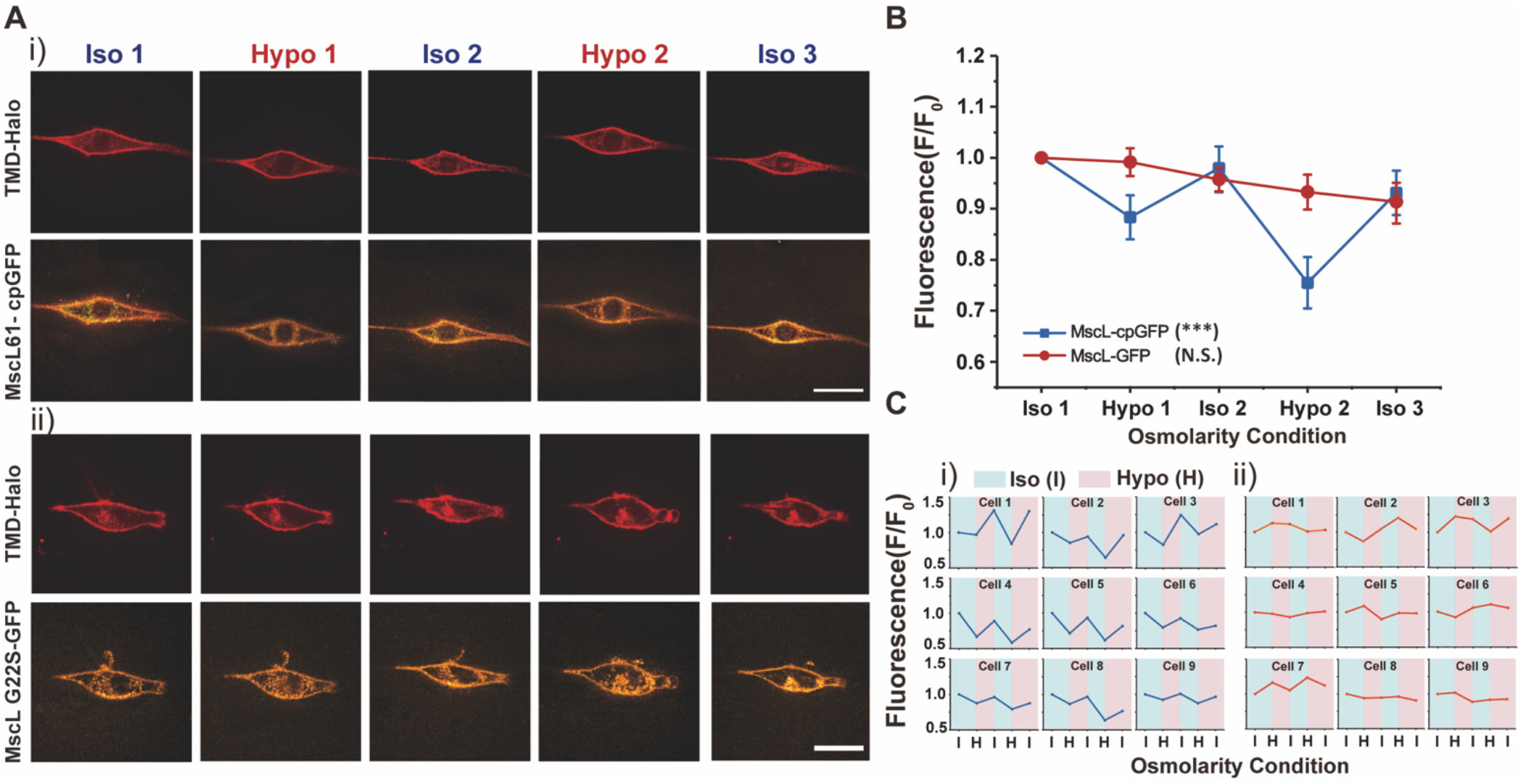
NIH3T3 cells transfected with the MscL tension biosensor in response to cyclic pressure (A) (i) Confocal images showing the membrane localization of MscL tension biosensor in NIH3T3 cells in response to the cyclic pressure test, which was carried out by alternating between iso-osmotic and hypo-osmotic conditions repeatedly. Transfected cells were cultured in iso-osmotic condition for 2 days. DI water was first added to the cell culture media to create hypo-osmotic environments and then 10X PBS solution was added back to the media to increase the osmolarity for iso-osmotic environments. The osmolarity of the media is ~330 mOsm for iso-osmotic conditions and ~165 mOsm for hypo-osmotic conditions. Each confocal image was taken four minutes after the addition of DI water or PBS solution. (ii) The confocal images showing the NIH3T3 cells transfected with the MscL(G22S)-c-term-GFP construct as a control in response to the same cyclic pressure cycle test as mentioned in (i). (B) Normalized fluorescence intensities of the cell membranes of NIH3T3 cells transfected with the MscL tension biosensor and MscL(G22S)-c-term-GFP construct corresponding to the cyclic pressure cycle test. Nine cells were analyzed for each condition from three independent experiments. (C) (i) Fluorescence intensity traces of MscL(G22S)61-cpGFP on cell membranes of nine different NIH3T3 cells during the cyclic osmotic tests (ii) Fluorescence intensity traces of MscL(G22S)-c-term-GFP on cell membranes of nine different NIH3T3 cells as a control. Scale bars: 10 μm. The error bars denote standard error. ***: *p* < 0.001.

All our osmotic shock measurements so far have been conducted at 4 min after the onsets of osmotic shocks. Nonetheless, it is well appreciated that cells have the ability to accommodate and recover from rapid changes in cell volume during osmotic shocks to regulate membrane tension^28^. Therefore, we sought to correlate the cell recovery dynamics from osmotic shocks to the corresponding membrane fluorescence of the MscL tension biosensor. We performed cell volume measurements every 1 min after hypo-osmotic shocks (with ~165 mOsm osmotic difference between cells and medium) in the first 5 min and continue tracking at 10 min, 20 min and 30 min, respectively, by using 3D reconstruction of side view of cells analyzed with Limeseg (Figure 4A). In separate experiments we acquired MscL tension biosensor fluorescence under the same condition. We observed significant increase in cell volume (~61%) and decrease in tension biosensor fluorescence (~21%) in the first 2 min after exposure of cells to hypotonic medium, which is followed by gradual decrease in the average cell volume from 2 to 10 min. During this same period, the corresponding fluorescence of the tension biosensor increased from 2 to 10 min (Figure 4B), showing a positive correlation between cell volume and membrane tension changes during the dynamic cell recovery under hypotonic condition.

**Figure 4.**
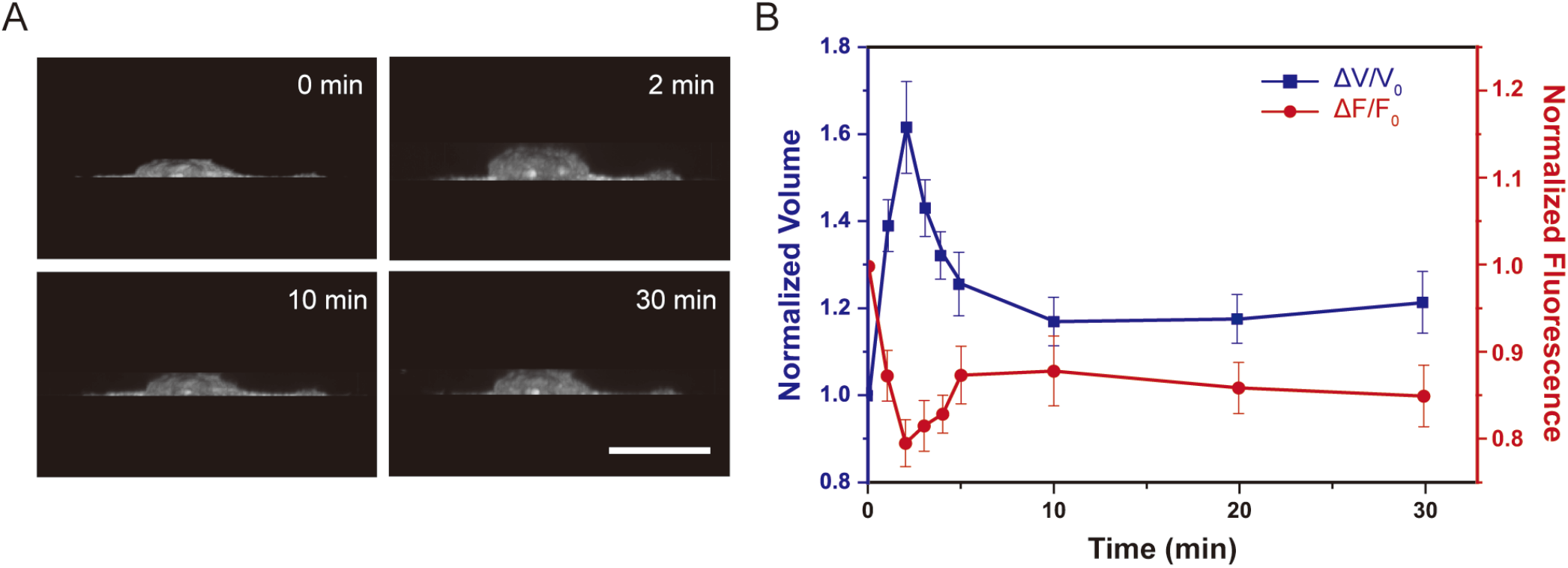
Volume and membrane fluorescence dynamics of NIH 3T3 cells expressing the MscL membrane tension biosensor (MscL(G22S)61-cpGFP) exposed to a hypotonic (~165mOsm) solution. (A) 3D reconstruction of cell volume using ImageJ 3D projection. (B)Normalized volume change and cpGFP fluorescence change under hypotonic shock of 165 mOsm. Confocal images were captured every 1 minute after osmotic shocks for the first 5 minutes followed by once every 5 minutes. Ten cells were analyzed for each condition from three independent experiments. Scale bars: 10 μm. The error bars denote standard error.

Finally, we used polydimethylsiloxane (PDMS) micropost arrays to further demonstrate the applicability of the MscL membrane tension sensor in cells under different spreading conditions. Micropost array technologies have been widely used to study the mechanosensitive behaviors of cells. Cells attached to fibronectin-coated microposts with different post heights, therefore stiffnesses, experience different degrees of cell spreading^29–31^. Cells attached to shorter microposts have greater membrane tension as they spread more on more rigid substrates. Conversely, longer microposts cause lower plasma membrane tension due to limited cell spreading on softer substrates. For our assay, fibronectin (50 μg/mL) was microcontact printed onto micropost arrays to create an adhesive surface. Human mesenchymal stem cells (hMSCs) co-transfected with the MscL tension biosensor and TMD-Halo were seeded at a density of around 1,500 cells/cm^2^ on micropost arrays with posts of 1.83 μm diameter and 2.3 or 6.4 μm height. We observed successful expression and membrane localization of the MscL tension biosensor, and cells seeded on short microposts exhibited lower cpGFP membrane fluorescence in compared to those attached to longer microposts (Figure 5A), consistent with the data from the osmotic pressure experiments in Figure 2Ai. We should note that cells were transfected prior to seeding them on short vs. long microposts, so differences in membrane tension biosensor fluorescence intensity in cells on short vs. long microposts could not have resulted from different expression levels. As expected, cells spread more on short microposts compared to long microposts, irrespective of the constructs expressed (Figure 5B). While cells expressing the MscL tension sensor responded to changes in cell spreading, with ~33% reduction in average of membrane fluorescent intensity from longer to shorter microposts, the control MscL(G22S)-c-term-GFP expressing cells showed no difference in GFP membrane fluorescence under the same conditions (Figure 5C).

**Figure 5.**
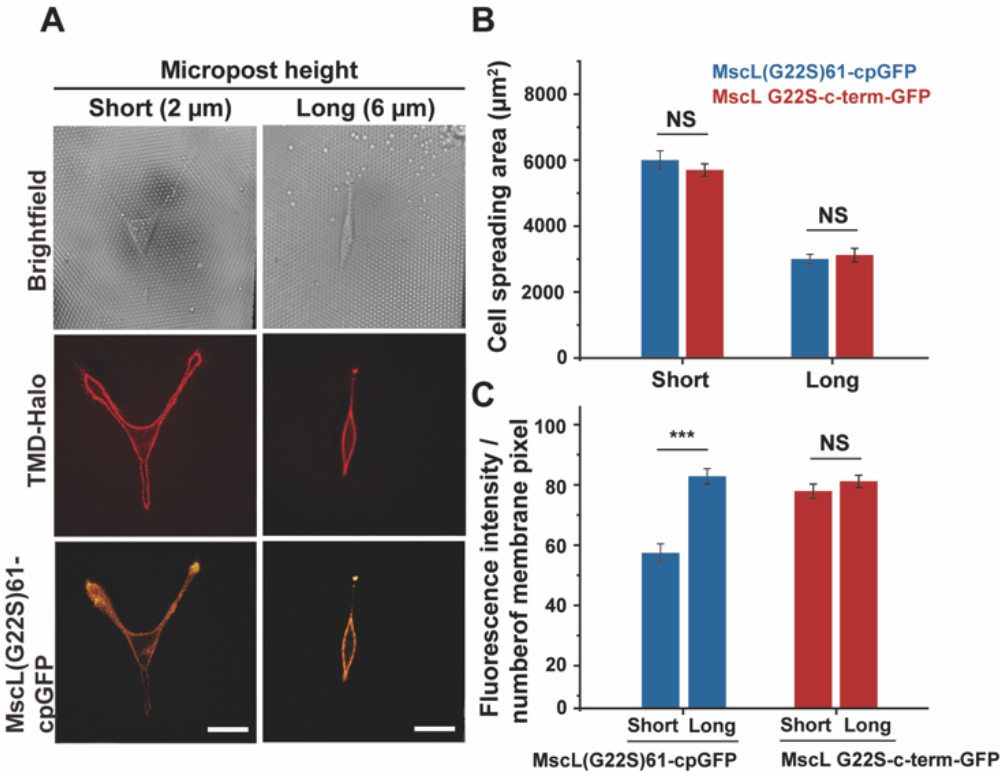
Fluorescence response of the MscL tension biosensor expressed in hMSCs attached to micropost arrays with different rigidity. (A) Confocal images showing the expression of the MscL tension biosensor in hMSCs spreading on short (left) or long (right) microposts. (B) Cell spreading areas of hMSCs on microposts with different heights. (C) The average fluorescence intensity of cpGFP or GFP signals localized to the cell membranes in response to different substrate rigidity. Nine cells were analyzed for each post height from three independent experiments. Scale bars: 50 μm. The error bars denote standard error. NS denotes not significant. *** represent *p* < 0.001.

Our study demonstrates that the MscL tension biosensor can be expressed efficiently on cell membranes and used in different cell types for membrane tension measurement. However, one drawback is the limitation that the biosensor only measures membrane tension in hypotonic conditions due to intact MscL structure under iso- and hypertonic conditions. Another potential concern is that MscL expression may exert major physiological effects on the cells and alter innate cell membrane tension and volume regulation, given MscL’s large opening. Previously, our lab has expressed MscL in mammalian cells and introduced membrane impermeable small molecules into cells following a 2 to 3-min hypo-osmotic shock^26^. We did not find MscL expression and activation had an adverse effect on cell viability. In another more recent work, we examined migration of MscL expressing cells in confining microfluidic channels and found that migration velocity was not affected once they entered the microfluidic channel, but MscL expressing cells were stuck more at the entrance compared to control cells^27^. Nevertheless, the effect of MscL activation on cell physiology should be further examined to ensure the use of the biosensor does not adversely impact cell physiology. In this context, it is perhaps worth speculating that the cpGFP inserted into the extracellular loop may reduce MscL conductance. Finally, it will be desirable to perform detailed analysis of membrane tension and fluorescence from the biosensor. Membrane tension response to hypo-osmotic shock likely varies significantly across cell types and such calibration, in each experimental system, would be necessary for quantitative assessment of membrane tension and would be the subject of future research. Given our goal was to establish a membrane tension biosensor based on MscL, we believe our current data support the use of this biosensor.

In summary, we successfully developed an optical membrane tension biosensor by repurposing MscL through insertion of cpGFP. As an OFF sensor, fluorescence of the MscL tension biosensor is negatively correlated to membrane tension. The MscL membrane tension biosensor is user friendly, thus offering a useful approach for detailed investigations of spatial and temporal dynamics of membrane tension, which will help advance research in cell mechanics, mechanotransduction, and cell signaling in a variety of cell types (e.g., mammalian, plant, and archaea) and subcellular organellar membranes including the membranes of the nucleus and the endoplasmic reticulum.

## ASSOCIATED CONTENT

### Supporting Information

The Supporting Information is available free of charge on the ACS Publications website.

Experimental details, micropost array fabrication, additional results, including figures and tables, data analysis and image processing (PDF)

## AUTHOR INFORMATION

### Author Contributions

A.P.L. and J.F. conceived the study. Y.Y.H., A.P.L. designed the experiments. Y.Y.H. performed the experiments. A.M.R. wrote the codes for image processing. Y.Y.H. and A.M.R. carried out image processing. Y.Y.H., A.M.R. and A.P.L. wrote the paper. All authors contributed to the manuscript revision and approved the final version.

## ACKNOWLEDGMENT

A construct with HaloTag-encapsulin was a kind gift from Tobias Giessen at the University of Michigan. We thank Taekjip Ha (Johns Hopkins University) for providing the pDisplay-SpyCatcher plasmid that contains Tf R transmembrane domain. We thank Robin Zhexuan Yan for helping with micropost array device fabrication. We thank Damon Hoff from the Single Molecule Analysis in Real Time Center at the University of Michigan for technical assistance. This work is supported by the National Institutes of Health (R21GM 134167 and R01EB030031). A.M.R.I. is partially supported by the Univ. Michigan Rackham Predoctoral Fellowship.

## For Table of Contents Only

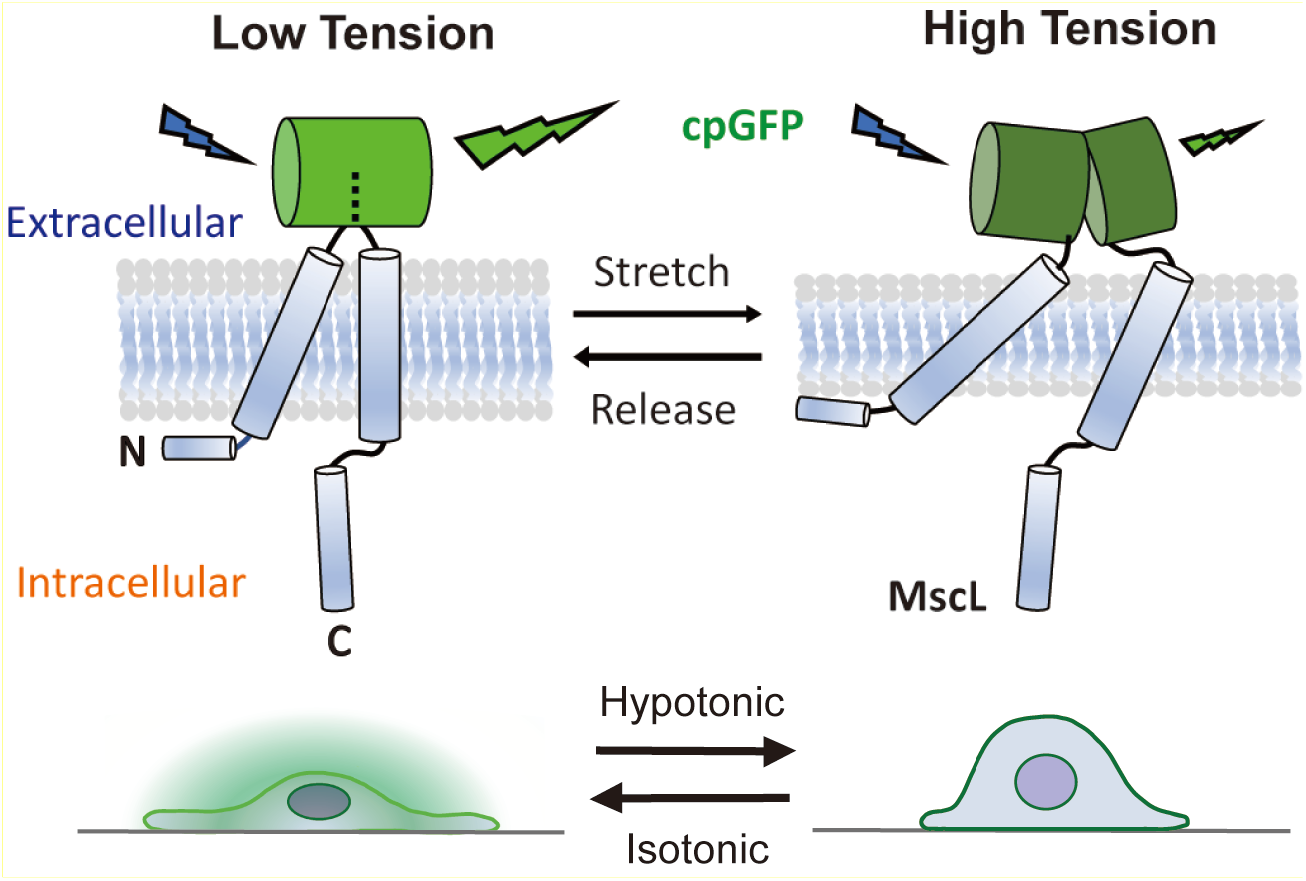

