## Supplementary materials for "Mechanosensitive channel-based optical membrane tension biosensor"

**Table of Contents**

Supporting information…………………………………………………………………………..S1

Experimental and analysis details………………………………………………………………..S3

Additional figures……………………………………………………………….................S13

Additional tables…………………………………………………………………………..S21

Reference………………………………………………………………………………….S28

### Experimental and analysis details

#### Materials and reagents

NIH3T3 cells were purchased from American Type Culture Collection. Human mesenchymal stem cells (hMSCs) and MSCGM™ Mesenchymal Stem Cell Growth Medium BulletKit™were purchased from Lonza Pharma and Biotech. Gibco Dulbecco's Modified Eagle Medium (DMEM), trypsin-EDTA (0.25%), and phosphate-buffered saline (PBS, 1X) were purchased from Thermo Fisher Scientific. Fetal bovine serum (FBS) was purchased from Gibco. Human fibronectin, Pluronic® F-127, and Trichloro (1H,1H,2H,2H-perfluorooctyl) silane were purchased from Sigma-Aldrich. SYLGARD™ 184 Silicone Elastomer Kit (polydimethylsiloxane (PDMS) base & curing agent) was purchased from Dow. All primers for cloning were purchased from Eurofins Scientific.

#### DNA constructs

pDisplay-SpyCatcher was a kind gift from Dr. Taekjip Ha, Johns Hopkins University. Linear HaloTag-encapsulin dsDNA was a kind gift from Dr. Tobias Giessen, University of Michigan. P70a-G-GECO was used in in our previous study[1]. pLVX-mCherry-P2A-MscL G22S-Puro was used in a separate previous study[2].

To generate MscL G22S-cpGFP construct as a membrane tension sensor for cell transfection, MscL G22S construct in pLVX-Puro was used as a template for Gibson assembly cloning. The primers used in the study are included in **Table S1**. First, cpGFP was amplified from P70a-G-GECO construct using the primers cpGFP – F and cpGFP – R with Phusion High-Fidelity DNA polymerase. Next, cpGFP was inserted into MscL G22S after amino acid 61 in the pLVX-mCherry-P2A-MscL G22S-Puro construct. To amplify the backbone, pLVX-mCherry-MscL G22S-Puro, primers MscL61 G22S-F and MscL61 G22S-R were used with Phusion High-

Fidelity DNA polymerase for PCR amplification. Afterwards, the resulting PCR products, cpGFP and pLVX-mCherry-P2A-MscL G22S-Puro were digested with DpnI for 1 hour at 37 °C and subsequently purified with the QIAquick Gel Extraction Kit, and then ligated together with homemade Gibson Master Mix (**Table S2**) to create pLVX-mCherry-P2A-MscL61 G22S-cpGFP-Puro construct.

MscL G22S -cpGFP construct with ER export signal sequence, TTTTGCTATGAAAATGAAGTT, was generated to increase MscL's membrane localization. pLVX-mCherry-P2A-MscL(G22S)61-cpGFP-Puro or pLVX-EGFP-P2A-MscL(G22S)61-cpGFP-Puro was used as a template respectively for Gibson assembly cloning. To encode ER export signal after MscL(G22S)61-cpGFP, ER export signal sequence was designed to be included within the primer ERexp – R. Combined with the primer ERexp – F, MscL61 G22S with cpGFP inserted and followed by the ER export signal sequence, referred to as MscL61 G22S-cpGFP-ERexp, was produced and amplified by PCR. The rest of the part of the template, which was pLVX-mCherry-P2A-Puro or pLVX-EGFP-P2A-Puro, was defined as the backbones and was amplified using the primers MscL61 G22S – cpGFP-F and MscL61 G22S – cpGFP-R. Again, the resulting PCR products were digested with DpnI, subsequently purified, and then ligated to create pLVX-mCherry-P2A-MscL(G22S)61-cpGFP-ERexp-Puro and pLVX-EGFP-P2A-MscL(G22S)61-cpGFP-ERexp-Puro construct.

To remove mCherry-P2A from the pLVX-mCherry-P2A-MscL(G22S)61-cpGFP-Puro construct, or to remove EGFP-P2A from the pLVX-EGFP-P2A-MscL(G22S)61-cpGFP-Puro construct, primers Del-mCherry/EGFP-MscL-F and Del-mCherry/EGFP- MscL-R were used to amplify the vector (pLVX – Puro construct) without mCherry or EGFP, which contains MscL(G22S) 61 with cpGFP and ER export signal. The resulting PCR products were digested with DpnI, subsequently purified, and then ligated to create pLVX -MscL(G22S)61-cpGFP-ERexp-Puro, which is the final version of our MscL membrane tension sensor. For the control, cpGFP was removed and GFP was fused to the C terminus of MscL G22S to generate pLVX-MscL G22S-c-term-GFP-ERexp-Puro construct. To replace MscL(G22S)61-cpGFP with MscL G22S-GFP in the pLVX-Puro vector, primers MscL-GFP-F and MscL-GFP-R were used. Primers pLVX-Puro-F and pLVX-Puro-R were also used to amplify pLVX-Puro vector as the backbone. The resulting PCR products were digested with DpnI, subsequently purified, and then ligated to create pLVX-MscL(G22S)61-GFP-ERexp-Puro construct.

pLVX-MscL G22S-c-term-cpGFP-ERexp-Puro, where cpGFP is fused to the C terminus of MscL G22S, and pLVX-Glu70-ERexp-cpGFP, where cpGFP is inserted into the extracellular domain of the glutamine transporter GluR0 after amino acid 70, were designed as other controls. To generate pLVX-MscL G22S-c-term-cpGFP-ERexp-Puro construct, primers Control-cpGFP-F and Control-cpGFP-R were used to amplify the insert fragment MscL G22S-c-term-cpGFP-ERexp, which is purchased from Twist Bioscience. For pLVX-Glu70-ERexp-cpGFP construct, primers Glu-F and Glu-R (same as Control-cpGFP-R) were used to amplify the insert fragment Glu70-ERexp, which is purchased from Twist Bioscience. Primers pLVX-Puro-F and pLVX-Puro-R were used to amplify pLVX-Puro vector as the backbone for both control constructs.

To label cell membranes, TMD-HaloTag construct was generated by encoding HaloTag, into the transmembrane domain of transferrin receptor (TfR) in pDisplay-SpyCatcher construct, a gift from Dr. Taekjip Ha at Johns Hopkins University, by Gibson assembly cloning. The linear DNA construct HaloTag-encapsulin with HaloTag fused to the *T. maritima* encapsulin capsid is a gift from Dr. Tobias Giessen at the University of Michigan. Initially, HaloTag was amplified from HaloTag-encapsulin using the primers HaloTag – F and HaloTag – R. To localize HaloTag to the cell membrane, it is fused to the transmembrane domain of TfR by replacing sfGFP with HaloTag in the pDisplay-SpyCatcher construct. To remove sfGFP from pDisplay-SpyCatcher construct as the backbone, primers xCatch-F and xCatch-R were used for PCR amplification. The resulting PCR products, HaloTag and pDisplay-SpyCatcher without sfGFP were digested with DpnI, subsequently purified, and ligated to create TMD-HaloTag construct. All constructs created in this study are summarized in **Table S3**.

**Cell culture and transfection**

NIH3T3 cells were cultured in 2 ml of growth medium (DMEM supplemented with 2 mM L-glutamine, 1 mM sodium pyruvate, and 10% fetal bovine serum (Gibco) and seeded onto 35 mm glass-bottom dishes (MatTek) for 2 days before transfection. For transfection, cells at 50% confluency were co-transfected with the desired DNA constructs (~500 ng/µl), MscL tension sensors version 1 or 2 (pLVX-MscL(G22S)61-cpGFP-Puro or pLVX-MscL(G22S)61-cpGFP-ERexp-Puro), or control plasmids (pLVX-GFP-P2A-MscL G22S-ERexp-Puro, pLVX-MscL G22S-c-term-GFP-ERexp-Puro, pLVX-MscL G22S-c-term-cpGFP-ERexp-Puro, pLVX-Glu70-cpGFP-ERexp-Puro) along with TMD-HaloTag, using Lipofectamine 3000 transfection reagent (Invitrogen) for 2 days before they were subjected to osmotic pressure experiments. Constructs used for transfections are listed in **Table S4**.

hMSCs were cultured in 2 ml of growth medium and seeded onto 6-well culture plate for 2 days before transfection. For transfection, cells at 70% confluency were co-transfected with the desired DNA constructs (~500 ng/µl), MscL tension sensors (pLVX-MscL(G22S)61-cpGFP-ERexp-Puro) or control plasmids (pLVX-MscL G22S-c-term-GFP-ERexp-Puro or pLVX-GFP-P2A-MscL G22S- ERexp-Puro) along with TMD-HaloTag, using Lipofectamine 3000 transfection reagent for 1 day. The transfected cells were then trypsinized and reseeded on the micropost arrays at a density of 1500 cells/cm$^2$. hMSCs were allowed to attach and spread on the micropost for at least 2 hours before imaging.

**Membrane labeling with HaloTag fluorescent ligand**

10 µl of 1 mg/ml JF594i-HaloTag ligand (obtained from Janelia Research Campus) was added to the 35 mm glass bottom dish with or without micropost arrays containing 2 ml of growth medium with cells transfected with TMD-HaloTag. The plate was incubated for 10 min at 37°C before washing with 2 ml of 1X PBS, pH 7.4 three times to remove the residual ligands. Afterwards, 1.5 ml of the fresh culture medium (DMEM supplemented with 10% fetal bovine serum) was added to the plate and incubated for 20 min at 37°C with 5% $CO_2$ before imaging.

**Application of hypo-osmotic/hyper-osmotic shock and pressure cycle test**

The osmolarity of DMEM supplemented with 10% FBS was measured by the vapor-pressured osmometer (ELITech Group) to be around 330 mOsm. In the osmotic pressure test, the osmolarity of the growth medium in the dish was progressively decreased from 330 to 108 mOsm or increased from 330 to 552 mOsm (step by step) by adding water or 10X PBS in order to apply increasing or decreasingosmotic shock to the cells, which stay in the desired osmotic condition for 4 minutes before image acquisition and the next addition of water/ 10X PBS. To achieve a targeted osmolarity, different amount of ultrapure water/10X PBS was added to 1 ml growth medium as shown in **Table S5.**

For the pressure cycle test, the osmolarity of the growth medium was alternated between ~330 mOsm (iso-osmotic condition) and ~165 mOsm (hypo-osmotic condition). Cells were subjected to two times of hypo-osmotic shocks in between three times of iso-osmotic conditions.

First, 1000 µl ultrapure water was added to 1 ml growth medium to create hypo-osmotic solution, and then 127 µl 10X PBS solution (~2930 mOsm) was added back to the growth medium to increase the osmolarity back to 330 mOsm for creating iso-osmotic environment. To apply second osmotic shock to cells, 2127 µl ultrapure water was added to the medium. For the last step, addition of 270 µl 10X PBS solution returned the sample to iso-osmotic condition again. The stepwise procedures for the amount of PBS solution and water added to the growth medium is summarized in **Table S6**.

### Micropost array fabrication

<u>Replica molding of micropost arrays</u>

Soft lithography was used to fabricate patterned PDMS stamps from silicon molds fabricated using photolithography and deep reactive-ion etching, as described previously[3]. First, the PDMS base and crosslinker were mixed at a 10:1 ratio and degassed under vacuum for 0.5 hr to remove bubbles. The uncured PDMS mixture was poured over the SU-8 silicon master in an aluminum boat before baking at 110 °C for 20 minutes in the oven. The partially cured PDMS negative molds were then peeled away from silicon master slowly, treated with oxygen plasma for 90 seconds to activate the surface, and left overnight to silanize in a desiccator using trichlorosilane. For positive casting, a droplet of 10:1 w/w PDMS was placed onto each PDMS mold and topped by a plasma-treated 18 mm-diameter round glass coverslip. The assembled PDMS molds were then placed in an oven at 110°C for 20 hrs to fully cure. After curing, the negative PDMS molds were slowly peeled away from the glass coverslips and inspected for collapsed posts. To restore micropost integrity, the patterned coverslips were immersed in 100% ethanol, sonicated for 1 minute, and subjected to dry-release in a critical point drier (Tousimis

Samdri-PVT-3D). The micropost array-containing coverslips were then adhered onto bottomless 12-well plates with the use of 10:1 w/w PDMS.

PDMS stamps for microcontact printing

Microcontact printing was carried out on the micropost array to control the shape or spreading of cells. First, stamps were generated by pouring PDMS (10:1 mixture of PDMS base and crosslinker) into a 150 mm Petri dish and cured at 65 °C for at least 2 hours. The PDMS was then peeled from the wafer, cut into blocks with dimensions comparable to the micropost arrays, and coated with 100 µl of 50 µg/mL fibronectin 1 hour. Finally, the stamps were washed with DI water and dried with nitrogen gas. The micropost substrates were ozone treated for 7 minutes to activate the surface for stamping. The fibronectin-coated stamps were gently lowered and placed in conformal contact with the micropost arrays, lightly tapping the top with tweezers while being careful to avoid post collapse. The stamps were then carefully removed with tweezers. The microposts were then sterilized using submersion in 100% ethanol followed by submersion in 70% ethanol for 15 seconds each. Finally, the substrate was washed with DI water three times. After washing, the micropost substrates were submerged in 0.2% Pluronic F-127 (Sigma-Aldrich) for 30 minutes in order to prevent the adsorption of additional proteins to areas outside of the stamped pattern. The substrates were then rinsed with DI water three times and stored in DI water at 4°C for up to one week.

Cell seeding

To seed hMSCs onto the microposts, we first replaced the DI water in the wells with mesenchymal stem cell growth medium. hMSCs were washed with Dulbecco's phosphate buffered saline and trypsinized with 0.25% trypsin-EDTA at 37°C for 5 minutes. The cell solution was then centrifuged at 1.5k rpm for 5 minutes. After removing trypsin-EDTA, the cells were resuspended in the growth medium and a density of approximately 1,500 cells/cm$^2$ was pipetted into each micropost array well. The dish was then placed in the incubator for at least 2 hours to allow cells to adhere and spread onto the adhesive islands of the micropost arrays.

**Fluorescence imaging**

All images were acquired using an oil immersion 60×/1.4 NA Plan-Apochromat objective with an Olympus IX-81 inverted fluorescence microscope (Olympus, Japan) controlled by MetaMorph software (Molecular Devices) equipped with a CSU-X1 spinning disk confocal head (Yokogawa, Japan), AOTF-controlled solid-state lasers (Andor, Ireland) or via a custom controller, and an iXON3 EMCCD camera (Andor). Images of cpGFP/GFP fluorescence and HaloTag fluorescence were acquired with 488 nm laser excitation at an exposure time of 500 ms and with 561 nm laser excitation at an exposure time of 200 ms, respectively. Each acquired image contained 1-3 cells. For an individual experiment, at least three images with nine cells in total were taken at different locations across a well. Three independent repeats were carried out for each experimental condition. Cells were stained with JF594i-HaloTag ligand for 10 minutes at 37°C prior to imaging.

**Data analysis and image processing**

Images were processed in Python with the use of OpenCV. The membrane signal was isolated using a multi-step background subtraction pipeline. For the first background subtraction step, a binary threshold (cv2.threshold) was applied to the JF594i-HaloTag signal (561 channel) to obtain a background mask (Figure S1 B). Background noise was eliminated by finding the connected components (cv2.connectedComponentsWithStats) and filtering them out by size. The mask was further smoothed using closing (cv2.MORPH_CLOSE) and opening (cv2.MORPH_OPEN) operations. Following this step, the component filtering process was repeated. This background mask was then used as a mask to eliminate background noise (Figure S1 C). For the second background subtraction step, this output image with reduced background noise was subjected to adaptive gaussian thresholding (Figure S1 D). This new binary image was then used as a mask to accurately capture the membrane signal from the 488 channel reporting the tension response (Figure S1 F). Total pixel intensity was measured and compared between different osmotic pressures. For the micropost array assay, average membrane signal of the membrane tension biosensor was measured for each condition. For this process, the membrane signal was isolated using the previously explained process, and the total signal intensity was divided by the number of pixels in the membrane to account for different cell sizes.

To better visualize the membrane cpGFP or GFP signal expressed in cells, the confocal images were adjusted using the lookup table function with orange hot selection built in Image J. However, the raw data captured by the confocal microscope was used for all data analysis and fluorescence calculation.

For fluorescence intensity of membrane pixels corresponding to osmolarity in Figure 2, and cell spreading areas and average fluorescent intensity of membrane pixels in Figure 4, the statistical analyses were verified by one-way ANOVA test to determine the $p$ values. The convention of $p$ values of *: $p < 0.05$; **: $p < 0.01$; ***: $p < 0.001$. $p < 0.05$ was considered statistically significant. Every experiment was performed three times with data collected from nine cells in each replicate.

For fluorescence intensity of membrane pixels analysis with cyclic osmolarity changes in Figure 3, statistical analysis was performed using a two-tailed $t$-test with a significance level of

0.05. The quantitative data was compared/analyzed within individual groups (within cells transfected with the MscL membrane tension sensors or control cells) between iso- and hypo-osmotic conditions. $p < 0.05$; **: $p < 0.01$; ***: $p < 0.001$. $p < 0.05$ was considered statistically significant. Every experiment was performed three times with data collected from nine cells in each replicate.

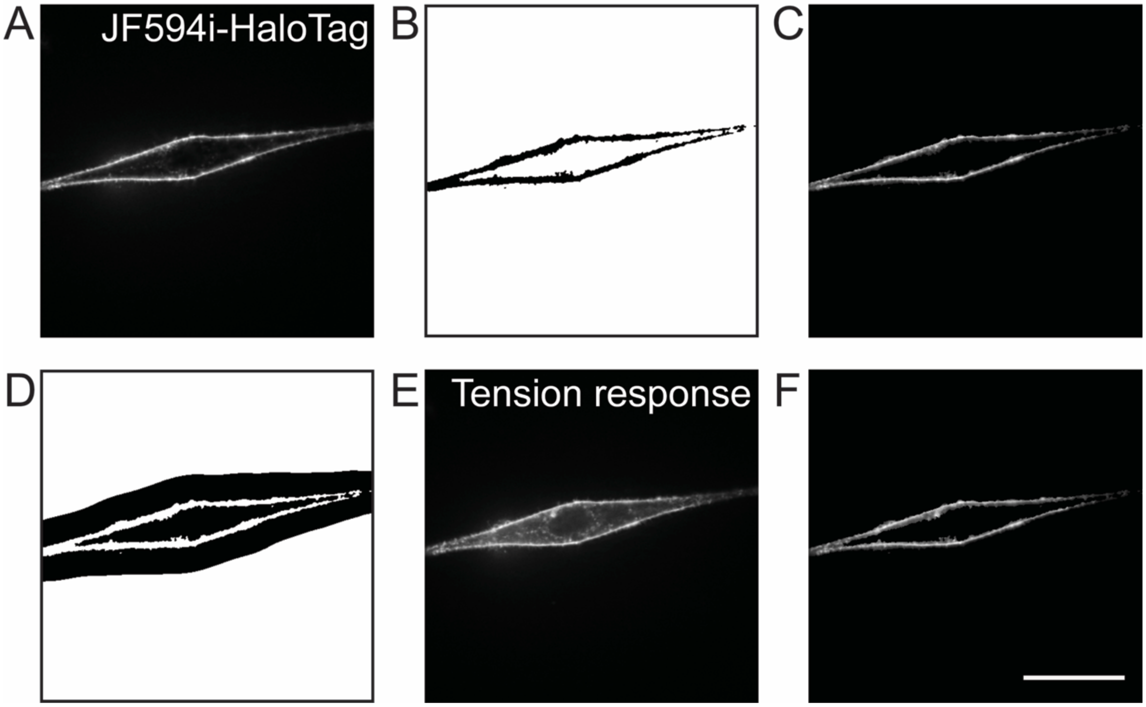


**Figure S1**. Multi-step background subtraction pipeline for tension signal measurement. (A) JF594i-HaloTag signal (561 channel). (B) Mask created by the application of a binary threshold to the JF594i-HaloTag signal. (C) JF594i-HaloTag signal with applied background mask. (D) Mask created from subjecting masked JF594i-HaloTag signal to adaptive Gaussian thresholding. (E) Tension response (488 channel). (F) Tension response with applied background mask resulting from adaptive Gaussian thresholding. Scale bar: 10 µm.

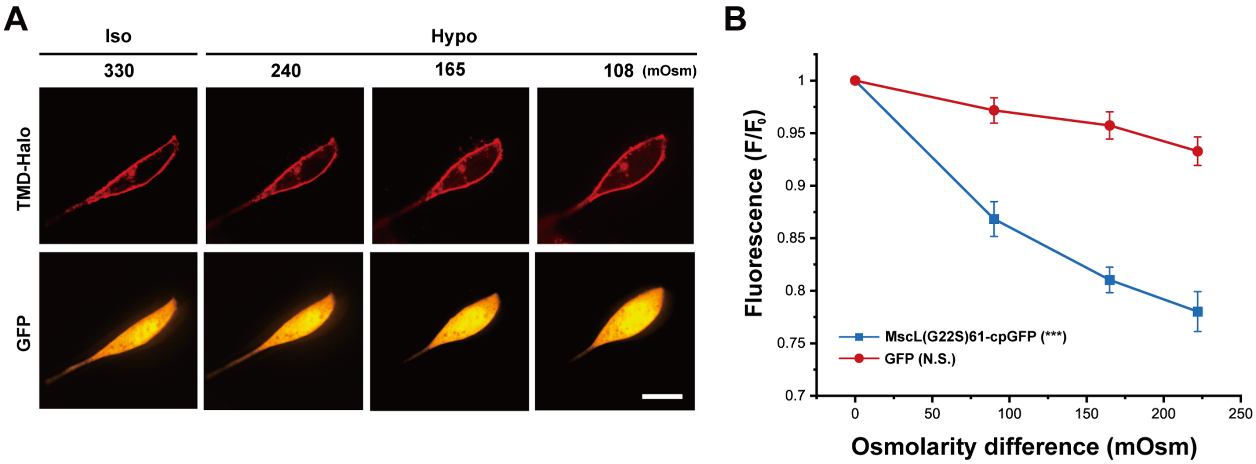


**Figure S2**. NIH3T3 cells expressing MscL G22S and GFP in response to hypo-osmotic pressure. (A) Confocal images showing the NIH3T3 cells transfected with the GFP-P2A-MscL(G22S) construct as a control in response to different osmotic shocks. Transfected cells were cultured in iso-osmotic condition for 2 days and then DI water was sequentially added to the cell culture media to create increasing hypo-osmotic environments. Each image was taken four minutes after the addition of DI water. (B) Normalized fluorescence intensities (MscL-cpGFP or GFP) of the cell membranes of NIH3T3 cells transfected with the MscL tension biosensor or GFP-P2A-MscL(G22S) under different osmotic conditions corresponding to the experiment mentioned in (A). Nine cells were analyzed for each condition from three independent experiments. Scale bars: 10 μm. ***: $p < 0.001$.

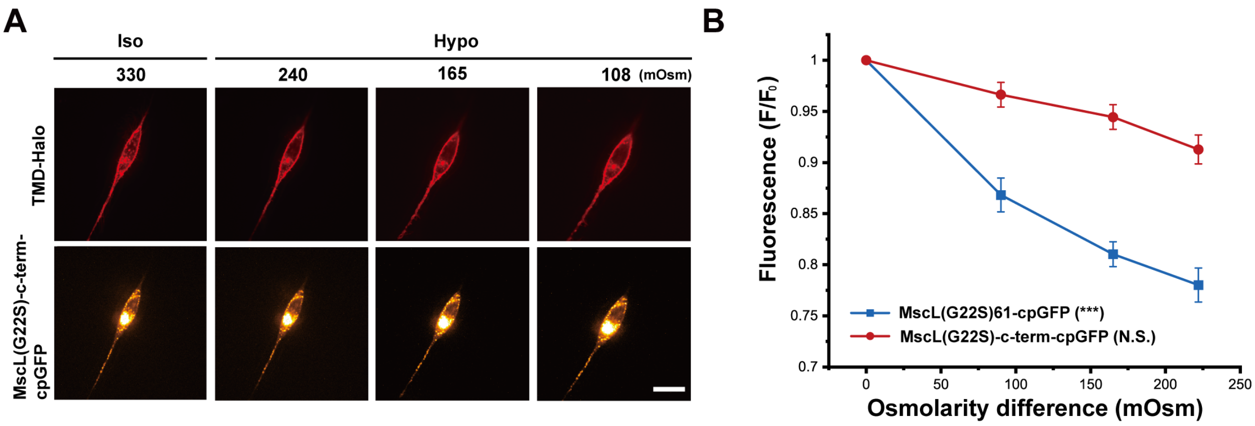


**Figure S3**. NIH3T3 cells expressing cpGFP fused to the c-terminus of MscL G22S in response to hypo-osmotic pressure. (A) Confocal images showing the NIH3T3 cells transfected with MscL(G22S)-c-term-cpGFP construct as a control in response to different osmotic shocks. Transfected cells were cultured in iso-osmotic condition for 2 days and then DI water was sequentially added to the cell culture media to create increasing hypo-osmotic environments. Each image was taken four minutes after the addition of DI water. (B) Normalized fluorescence intensities of the cell membranes of NIH3T3 cells transfected with the MscL tension biosensor or MscL(G22S)-c-term-cpGFP under different osmotic conditions corresponding to the experiment mentioned in (A). Nine cells were analyzed for each condition from three independent experiments. Scale bars: 10 µm. ***: $p < 0.001$.

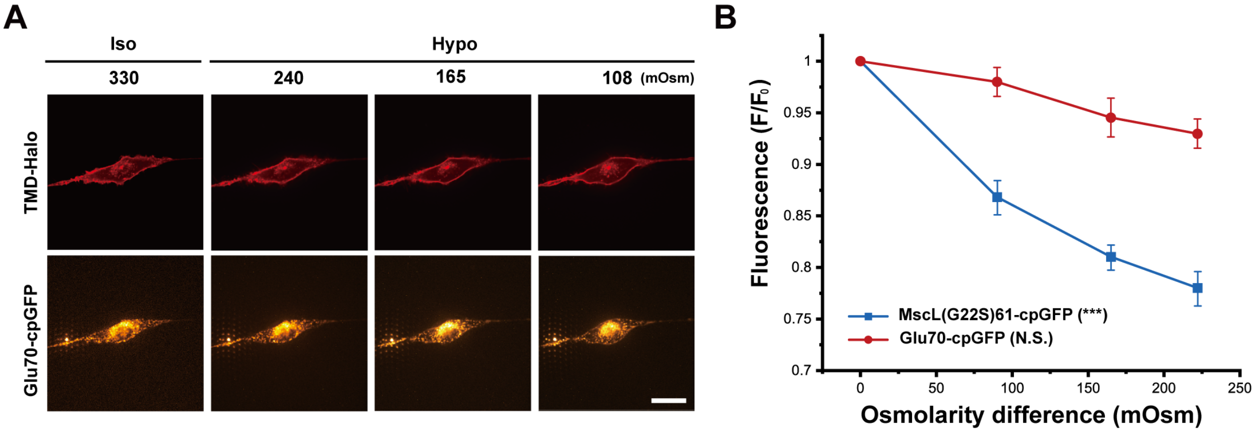


**Figure S4**. NIH3T3 cells expressing GluR0 with cpGFP inserted into its extracellular loop in response to hypo-osmotic pressure. (A) Confocal images showing the NIH3T3 cells transfected with the Glu70-cpGFP construct as a control in response to increasing osmotic downshock. Transfected cells were cultured in iso-osmotic condition for 2 days and then DI water was sequentially added to the cell culture media to create increasing hypo-osmotic environments. Each confocal image was taken four minutes after the addition of DI water. (B) Normalized fluorescence intensities of the cell membranes of NIH3T3 cells transfected with the MscL tension biosensor or Glu70-cpGFP construct corresponding to the experiment mentioned in (A). Nine cells were analyzed from three independent experiments. Scale bars: 10 µm. ***: $p < 0.001$.

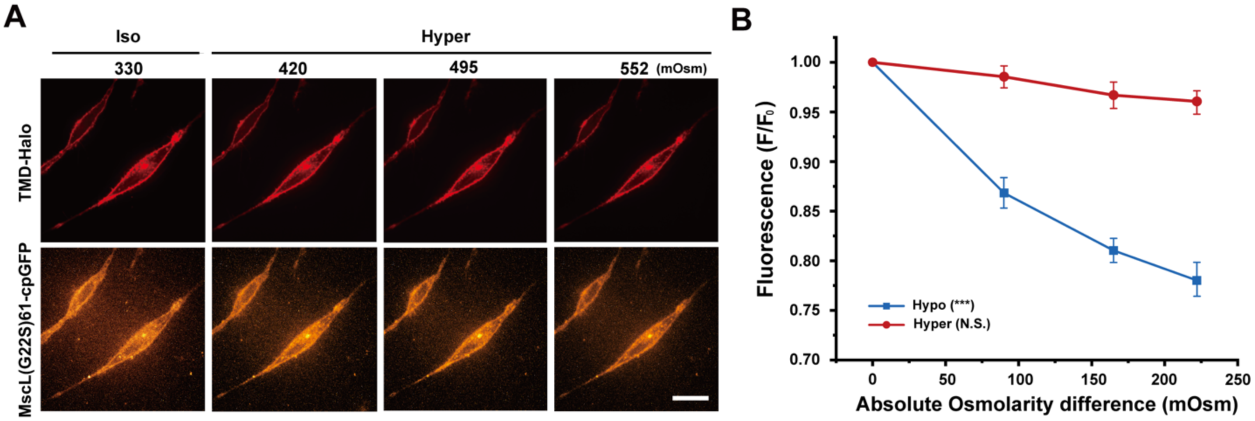


**Figure S5**. NIH3T3 cells expressing MscL(G22S)61-cpGFP in response to hyper-osmotic pressure. (A) Confocal images showing the NIH3T3 cells transfected with the MscL(G22S)61-cpGFP construct in response to different hyper-osmotic shocks. Transfected cells were cultured in iso-osmotic condition for 2 days and then 10X PBS solution was sequentially added to the cell culture media to create increasing hyper-osmotic environments. Each image was taken four minutes after the addition of DI water. (B) Normalized fluorescence intensities (hypo- or hyper-osmotic conditions) of the cell membranes of NIH3T3 cells transfected with the MscL tension biosensor under different osmotic conditions corresponding to the experiment mentioned in (A). Hypo-osmotic condition was described in Figure S4. Osmotic difference is higher external osmolarity for hyper-osmotic condition and lower external osmolarity for hypo-osmotic condition. Nine cells were analyzed for each condition from three independent experiments. Scale bars: 10 µm. ***: $p < 0.001$.

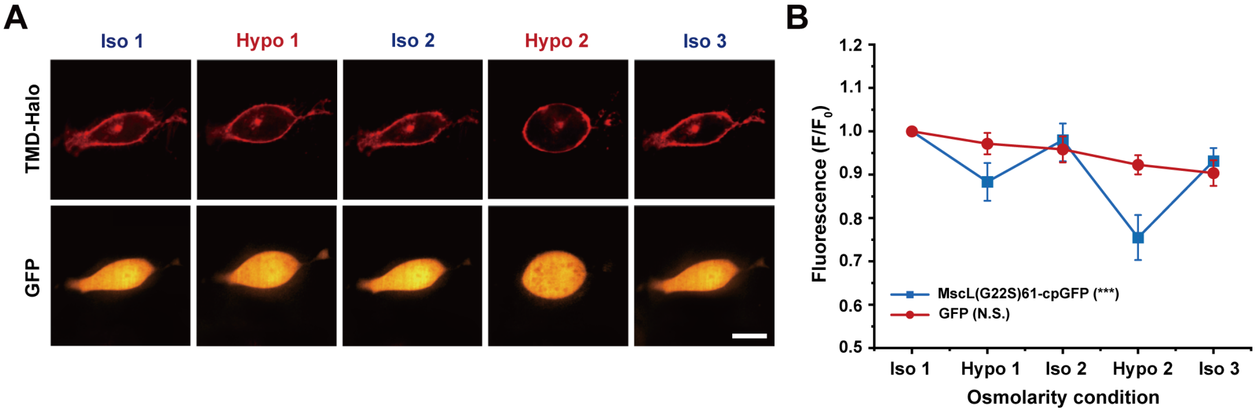


**Figure S6**. NIH3T3 cells expressing GFP in response to cyclic pressure. (A) Confocal images showing the NIH3T3 cells transfected with the GFP-P2A-MscL(G22S) construct as a control in response to the cyclic pressure test, which was carried out by alternating between iso-osmotic and hypo-osmotic conditions. Transfected cells were cultured in iso-osmotic condition for 2 days. DI water was first added to the cell culture media to create hypo-osmotic environments and then 10X PBS solution was added back to the media to increase the osmolarity back to iso-osmotic environments. The osmolarity of the media is ~330 mOsm for iso-osmotic conditions and ~165 mOsm for hypo-osmotic conditions. Each confocal image was taken four minutes after the addition of DI water or PBS solution. (B) Normalized fluorescence intensities of the cell membranes of NIH3T3 cells transfected with the MscL tension biosensor and GFP-P2A-MscL(G22S) construct corresponding to the cyclic pressure cycle test. Nine cells were analyzed from three independent experiments. Scale bars: 10 µm. ***: $p < 0.001$.

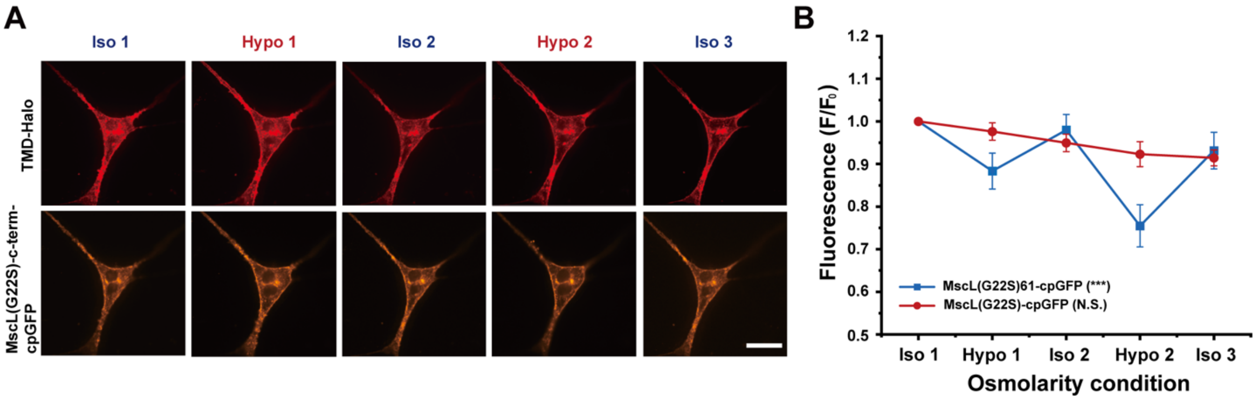


**Figure S7**. NIH3T3 cells expressing cpGFP connected to MscL G22S in response to cyclic pressure. (A) Confocal images showing the NIH3T3 cells transfected with the MscL(G22S)-c-term-cpGFP construct as a control in response to the cyclic pressure test, which was carried out by alternating between iso-osmotic and hypo-osmotic conditions. Transfected cells were cultured in iso-osmotic condition for 2 days. DI water was first added to the cell culture media to create hypo-osmotic environments and then 10X PBS solution was added back to the media to increase the osmolarity back to iso-osmotic environments. The osmolarity of the media is ~330 mOsm for iso-osmotic conditions and ~165 mOsm for hypo-osmotic conditions. Each confocal image was taken four minutes after the addition of DI water or PBS solution. (B) Normalized fluorescence intensities of the cell membranes of NIH3T3 cells transfected with the MscL tension biosensor and MscL(G22S)-c-term-cpGFP construct corresponding to the cyclic pressure test. Nine cells were analyzed from three independent experiments. Scale bars: 10 μm. ***: $p < 0.001$.

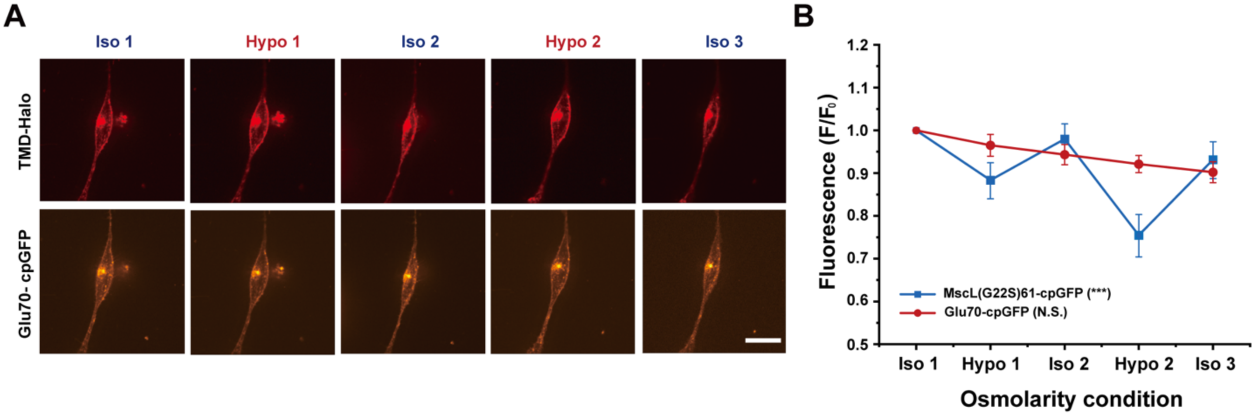


**Figure S8**. NIH3T3 cells expressing GluR0 with cpGFP inserted into its extracellular loop in response to cyclic pressure. (A) Confocal images showing the NIH3T3 cells transfected with the Glu70-cpGFP construct as a control in response to the cyclic pressure test, which was carried out by alternating between iso-osmotic and hypo-osmotic conditions. Transfected cells were cultured in iso-osmotic condition for 2 days. DI water was first added to the cell culture media to create hypo-osmotic environments and then 10X PBS solution was added back to the media to increase the osmolarity back to iso-osmotic environments. The osmolarity of the media is ~330 mOsm for iso-osmotic conditions and ~165 mOsm for hypo-osmotic conditions. Each confocal image was taken four minutes after the addition of DI water or PBS solution. (B) Normalized fluorescence intensities of the cell membranes of NIH3T3 cells transfected with the MscL tension biosensor and Glu70-cpGFP construct corresponding to the cyclic pressure cycle test. Nine cells were analyzed in each independent experiment with three in total. Scale bars: 10 µm. ***: $p < 0.001$.

**Table S1. List of primers used in this study. (Extended)**

| | |
|---|---|
| **cpGFP – F** | GATTTTAAACAGTTTGCTGTCACGAGCAAGGGCGAGGAGCTG |
| **cpGFP – R** | GATATCCCCCTGCGCATCGCGTAGGTTGTACTCCAGCTTGTGCCCCAG |
| **MscL61 G22S-F** | GCACAAGCTGGAGTACAACCTACGCGATGCGCAGGGGGATATC |
| **MscL61 G22S -R** | CAGCTCCTCGCCCTTGCTCGTGACAGCAAACTGTTTAAAATCGATCCCG |
| **ERexp – F** | CATGGACGAGCTGTACAAGGGAAGCGGAGCTACTAACTTCAGCCTGCTGAAGC |
| **ERexp – R** | CTACCCGGTAGAATTATCTAGATTAAACTTCATTTTCATAGCAAAAAGAGCGG |
| **MscL61 G22S – cpGFP-F** | TAATCTAGATAATTCTACCGGGTAGGGGAG |
| **MscL61 G22S – cpGFP-R** | TCCGCTTCCCTTGTACAGCTCG |
| **MscL-GFP-F** | CAGATCTCGAGCTCAAGCTTCGAATTCATGAGCATTATTAAAGAATTTCGCG |
| **MscL-GFP-R** | CTCCCCTACCCGGTAGAATTATCTAGATTACACTTCGTTTTCATAGCAAAAGCT |

| | |
|---|---|
| **Control-cpGFP-F** | CAGATCTCGAGCTCAAGCTTCGAATTCATGAGCATTATTAAAGA ATTTCGCGAATTTGCG |
| **Control-cpGFP-R/ Glu-R** | GCGCCTCCCCTACCCGGTAGAATTATCTAGATTACACTTCGTTT TCATAGCAAAAGCTGC |
| **Glu-F** | GCTACCGGACTCAGATCTCGAGCTCAAGCTTCGAATTCATGAGC GGAATTGGCCTTCTGATC |
| **pLVX-Puro-F** | TAATCTAGATAATTCTACCGGGTAGG |
| **pLVX-Puro-R** | GAATTCGAAGCTTGAGCTCGAG |
| **HaloTag – F** | ATCGGCACGGGCTTTCCGTTTG |
| **HaloTag – R** | TTCCAGCCCGGAGATCTCCAGTG |
| **xCatch-F** | CTGGAGATCTCCGGGCTGGAAGAATTCCTCGAGGCGGCCGC |
| **xCatch-R** | CAAACGGAAAGCCCGTGCCGATCCCGGATCCTCCGCTTCCATA G |
| **Del-mCherry/EGFP-MscL-F** | ctaccggactcagatctcgagCTCAAGCTTCgaattcATGAGCATTATTAAAGA ATTTCGCGAATTTG |
| **Del-mCherry/EGFP-MscL-R** | CATCGCAAATTCGCGAAATTCTTTAATAATGCTCATGAATTCGA AGCTTGAGCTCGAGATCTG |

**Table S2. 2X Homemade Gibson master mix for 20 reactions.**

| | |
|---|---|
| 5X isothermal reaction buffer | 40 µl |
| 10U/µl T5 exonuclease | 0.1µl |
| 2U/µl Phusion polymerase | 2.5 µl |
| 40U/µl Taq DNA ligase | 20 µl |
| UltraPure DNase/RNase-Free Distilled Water | 37.4µl |
| Total | 100 µl |

Note: The mixture is split in 5 µl aliquots for each reaction and should be froze immediately with liquid nitrogen. The reactions can be stored at -80°*C* for up to 3 months.

**Table S3. List of plasmids used in this study.**

| Content | Description |
|---|---|
| pLVX-mCherry-P2A-MscL G22S-Puro | MscL G22S separated from mCherry |
| pLVX-GFP-P2A-MscL G22S-Puro | MscL G22S separated from GFP |
| pLVX-mCherry-P2A-MscL(G22S)61-cpGFP-ERexp-Puro | cpGFP inserted after amino acid 61 of MscL G22S, separated from mCherry, with addition of ER export signal at its C terminus |
| pLVX-MscL G22S-c-term-GFP-ERexp-Puro | GFP connected to MscL G22S |
| pLVX-MscL G22S-c-term-cpGFP-ERexp-Puro | cpGFP connected to MscL G22S |
| pLVX-Glu70-cpGFP-ERexp-Puro | cpGFP inserted after amino acid 70 of Glutamine transporter GluR01 |
| pLVX-MscL(G22S)61-cpGFP-Puro | Membrane tension sensor version 1 |
| pLVX-MscL(G22S)61-cpGFP-ERexp-Puro | Membrane tension sensor version 2 |

**Table S4. List of plasmids used for each experiment.**

| | **DNA construct** | **DNA amount in 2ml media** |
|---|---|---|
| **Version 1** | pLVX-MscL(G22S)61-cpGFP-Puro (1) + TMD-HaloTag (2) | (1) 1.25 μg<br>(2) 1.25 μg |
| **Version 2** | pLVX-MscL(G22S)61-cpGFP-ERexp-Puro (1) + TMD-HaloTag (2) | (1) 1.25 μg<br>(2) 1.25 μg |
| **Control 1** | pLVX-MscL G22S-c-term-GFP-ERexp-Puro (1) + TMD-HaloTag (2) | (1) 1.25 μg<br>(2) 1.25 μg |
| **Control 2** | pLVX-GFP-P2A-MscL G22S-ERexp-Puro (1) + TMD-HaloTag (2) | (1) 1.25 μg<br>(2) 1.25 μg |
| **Control 3** | pLVX-MscL G22S-c-term-cpGFP-ERexp-Puro (1) + TMD-HaloTag (2) | (1) 1.25 μg<br>(2) 1.25 μg |
| **Control 4** | pLVX-Glu70-cpGFP-ERexp-Puro (1) + TMD-HaloTag (2) | (1) 1.25 μg<br>(2) 1.25 μg |

**Table S5. Osmolarity of iso-isomotic, hypo-osmotic, and hyper-osmotic solutions added sequentially to an initial 1 mL volume for hypo-osmotic and hyper-osmotic pressure tests.**

| Osmolarity of medium (mOsm) | Medium condition | Addition of water or 10X PBS (µl) |
|---|---|---|
| 330 | Iso-osmotic | 0 |
| 240 | Hypo-osmotic | 375 (water) |
| 165 | Hypo-osmotic | 1000 (water) |
| 108 | Hypo-osmotic | 2055.5 (water) |
| 420 | Hyper-osmotic | 35.9 (10x PBS) |
| 495 | Hyper-osmotic | 67.8 (10x PBS) |
| 552 | Hyper-osmotic | 93.4 (10x PBS) |

**Table S6. The amount of water or 10X PBS added to an initial 1 mL volume for cyclic osmotic pressure test.**

| Osmolarity of medium (mOsm) | Medium condition | Addition of water (μl) | Addition of 10X PBS (μl) |
|---|---|---|---|
| 330 | Iso-osmotic | X | X |
| 165 | Hypo-osmotic | 1000 | X |
| 330 | Iso-osmotic | X | 127 |
| 165 | Hypo-osmotic | 2127 | X |
| 330 | Iso-osmotic | X | 270 |
